# Functional brain networks predicting sustained attention are not specific to perceptual modality

**DOI:** 10.1101/2024.05.15.594382

**Authors:** Anna Corriveau, Jin Ke, Hiroki Terashima, Hirohito M. Kondo, Monica D. Rosenberg

## Abstract

Sustained attention is essential for daily life and can be directed to information from different perceptual modalities including audition and vision. Recently, cognitive neuroscience has aimed to identify neural predictors of behavior that generalize across datasets. Prior work has shown strong generalization of models trained to predict individual differences in sustained attention performance from patterns of fMRI functional connectivity. However, it is an open question whether predictions of sustained attention are specific to the perceptual modality in which they are trained. In the current study we test whether connectome-based models predict performance on attention tasks performed in different modalities. We show first that a predefined network trained to predict adults’ *visual* sustained attention performance generalizes to predict *auditory* sustained attention performance in three independent datasets (N_1_=29, N_2_=60, N_3_=17). Next, we train new network models to predict performance on visual and auditory attention tasks separately. We find that functional networks are largely modality-general, with both model-unique and shared model features predicting sustained attention performance in independent datasets regardless of task modality. Results support the supposition that visual and auditory sustained attention rely on shared neural mechanisms and demonstrate robust generalizability of whole-brain functional network models of sustained attention.

## Introduction

The maintenance of attention to information over time is essential for daily activities such as driving to work or conversing with friends. Recent work in cognitive neuroscience has been aimed at identifying neural signatures of sustained attention ability with the goal of constructing models that generalize across people and datasets to predict individual differences in attention function. However, while sustained attention can be deployed to information from multiple perceptual (e.g., visual and auditory) modalities, it is an open question whether predictive models generalize across modality. For example, models trained to predict performance on visual sustained attention tasks may contain modality-specific features and therefore fail to generalize or generalize poorly to capture auditory sustained attention performance. Alternatively, features may capture modality-general aspects of attention and generalize broadly. Here, we construct and test the generalizability of models trained to predict sustained attention to visual and auditory stimuli from functional connections.

Identifying brain-based markers of cognition is beneficial both for understanding associations between functional brain organization and behavior and for developing predictive models. Network neuroscience provides a framework for the identification of interpretable neural signatures of cognition (Srivastava et al., 2022). One method, connectome-based predictive modeling (CPM), identifies functional connections, or edges, between brain regions whose strength is reliably associated with phenotypes across individuals (Finn et al., 2015; Rosenberg et al., 2016; Shen et al., 2017). This method has identified edge networks that predict sustained attention within and across samples of individuals (Rosenberg et al., 2016a, 2020; Yoo et al., 2022).

The utility of predictive models lies in their external validity, i.e., generalizability across independent datasets and contexts (Poldrack et al., 2020; Rosenberg & Finn, 2022; Scheinost et al., 2019). Successful generalizability across datasets ensures a model’s accuracy in the identification of relevant features as well as robustness to differences between samples. Predictive models of sustained attention constructed using CPM have demonstrated generalizability across datasets, as well as generalization to other attention tasks and attention-related symptoms (Rosenberg et al., 2016a; 2018, 2020; Yoo et al., 2022). Therefore, CPM successfully captures functional networks related to attention across contexts.

Sustained attention is often measured using tasks that require continuous vigilance for the detection or discrimination of rare stimuli (Mackworth, 1948; Langner and Eickhoff, 2013). While much work has investigated sustained attention to visual stimuli, attention can be deployed to other perceptual modalities, such as audition. Previous work has shown that the ability to sustain attention to visual and auditory information is reliable within individuals, suggesting that these abilities rely on shared cognitive mechanisms (Corriveau et al., 2024; Seli et al., 2011; Terashima et al., 2021). Work using electroencephalography and fMRI data has identified neural substrates underlying detection of both visual and auditory rare targets (Katayama & Polich, 1999; Kondo et al., 2023; Kim, 2014; Linden et al., 1999; Stevens et al., 2000), further supporting a modality-general neural basis of sustained attention. However, recent work shows that neurometabolites are differentially related to auditory and visual sustained attention (Kondo et al., 2023). Further, selective attention to visual or auditory information elicits both supramodal and modality-specific neural activation patterns (Smith et al., 2010; Stevens et al., 2000), suggesting that attending these modalities relies on distinct neural mechanisms as well. Therefore, the extent to which visual and auditory sustained attention networks are modality-specific may depend on the extent to which they rely on modality-specific neural mechanisms.

Here, we test the extent to which predictions from functional networks of sustained attention are biased by the perceptual modality in which they are trained. We find that a network previously defined to predict visual sustained attention predicts performance across datasets and modalities. Further, we show that models trained on auditory and visual tasks are highly generalizable across perceptual modalities. Even after the removal of features identified by both visual and auditory networks, i.e., modality-general features, models successfully predict cross-modality sustained attention ability. These results demonstrate that sustained attention relies on distributed patterns of connectivity. Additionally, they suggest that distributed patterns may be different between perceptual modality but still capture generalizable variance in sustained attention ability across datasets and modalities.

## Methods

### Dataset 1

The first dataset analyzed was described in detail by Kondo et al. (2022; 2023). This study was reviewed and approved by the Research Ethics and Safety Committees of Chukyo University and ATR-Promotions. Participants provided their written informed consent to participate in this study.

Participants (*N*=29, ages 20-35) were healthy Japanese adults who completed an fMRI scan consisting of two visual runs and two auditory runs of a gradual onset continuous performance task (*gradCPT*; Esterman et al., 2013; Rosenberg et al., 2013; Terashima et al., 2021). Data were collected using a 3T Magnetom Prisma MRI scanner (Siemens, Munich, Germany). Task runs were 400 seconds in length. A multiband echo-planar imaging (EPI) sequence was used to collect 205 volumes per task run with a repetition time (TR) of 2 seconds. Voxels were 2mm x 2mm x 2mm. The first five volumes of each run were discarded for data analysis.

The *gradCPT* was developed to measure sustained attention performance. In the task, stimuli gradually transition one into the next to avoid abrupt onsets. Visual runs of the *gradCPT* featured round, grayscale images of city (90%) and mountain (10%) scenes. Images transitioned from off to fully visible over 1.6 seconds such that a stimulus reached maximum visibility every 1.6 s. Images faded from peak visibility to off as presentation of the next image began. Participants were instructed to press a button for each city scene and withhold a button press for mountain scenes.

Stimuli for the auditory *gradCPT* were narrations from a foreign language database, excluding Japanese narrations to avoid presentation of a participant’s native language. Thus, participants used acoustic clues of the stimuli, rather than semantic clues, to judge the gender of voice streams. Narrations were performed by male (90%) and female (10%) voices and gradually transitioned from one to the next using sinusoidal ramps (Terashima et al., 2021) such that a voice reached maximum presentation every 1.6 seconds. Participants were instructed to press a button for male voices and withhold a button press for female voices.

Because stimuli faded from one to the next, key presses were assigned to trials in an iterative manner that first assigned unambiguous presses and then assigned more ambiguous ones. Unambiguous key presses occurring in a window from 70% presented to 40% disappeared were assigned to the current trial. Key presses that occurred outside the window were assigned to adjacent trials if no responses to those trials had been made. If no response was made to either adjacent trial, the key press was attributed to the closer trial. If either trial was an infrequent trial (mountain scene, female voice), the key press was assigned to the adjacent frequent trial. Sustained attention performance was quantified using a measure of sensitivity (*d’*) which is calculated as the normalized hit rate minus the normalized false alarm rate for each run.

### Dataset 2

The second dataset was collected at the MRI Research Center at the University of Chicago. Study procedures were approved by the Social and Behavioral Sciences Institutional Review Board at the University of Chicago. All participants provided their written informed consent prior to participation.

Participants (*N*=60) participated in at least one session of a two-session fMRI study collected approximately one week apart (mean time between sessions=10.88 days, SD=9.87 days). During both sessions, participants performed a 10-minute audio-visual continuous performance task (*avCPT*; Corriveau et al., 2024). Functional MRI data were collected on a 3T Philips Ingenia scanner. Volumes were collected using a multiband sequence with a repetition time of 1 second. Three volumes were removed from the start of each scan.

During the *avCPT*, streams of trial-unique images and sounds were presented simultaneously. Images were presented continuously for 1.2 seconds each whereas sounds were presented for 1 second with a 200 ms inter-trial interval to allow participants to distinguish individual sounds. Each task run was 500 trials in length. Images were indoor and outdoor scenes drawn from the SUN image database (Xiao et al., 2010). Sound stimuli were natural and manmade sounds drawn from online sound databases and cropped to be 1 s in length. Full details of stimulus curation procedures are described in Corriveau et al. (2024).

Before the task run, participants were instructed to make a button press to frequent stimuli (90%) from either the auditory or visual modality and to withhold a button press for infrequent stimuli (10%). They were told that the stimuli from the other modality were not relevant for the task. Over the two scan sessions, participants performed both the auditory and visual task and the order of task runs and frequent stimulus category was counterbalanced across participants.

For frequent trials, correct responses were trials in which participants responded before the onset of a new stimulus (within 1200 ms of trial start). However, to allow for the possibility of RTs longer than 1200 ms, we reassigned key presses for frequent trials which met the following criteria: (1) the participant made more than one key press for a trial with a frequent-category stimulus (2) the first key press was faster than 100 ms, and (3) no response was made to the previous frequent-category stimulus. In this case, the first key press was attributed to the previous trial. This reassignment of key presses is meant to more accurately account for accurate performance with slower response times. Press reassignment was rare in both visual (mean number of trials with presses reassigned=.548, SD=.861) and auditory sessions (mean number of trials with presses reassigned=3.81, SD=3.46), affecting less than .8% of the trials in each task. Therefore, this analytical decision has a negligible effect on results. Performance during the *avCPT* was calculated as sensitivity (*d’)*.

### Dataset 3

The final fMRI dataset analyzed was described in Walz et al. (2013) and shared on OpenNeuro (ds000116). This dataset contained runs from 17 adults (6 females, ages 20-40 years) who performed three auditory and three visual runs of an oddball task. Simultaneous fMRI and electroencephalography data were collected for the original study but only the fMRI data are analyzed here. Data were collected on a 3T Philips Achieva scanner. Each run consisted of 170 volumes collected with a 2 s TR. While the authors note that discarding of extra runs is unnecessary for the shared data, the first three volumes of each run were removed in keeping with a standard preprocessing pipeline. We do not expect this to affect the current results.

Task runs consisted of 125 stimuli presented for 200 ms with a variable inter-trial interval of 2-3 seconds. Participants were instructed to press a button for infrequent targets (20%) and could ignore standard stimuli (80%). In visual runs, standard trials consisted of a small green circle and target trials were the presentation of a large red circle. For auditory runs, the standard stimulus was a 390 Hz tone, whereas the target stimulus was a broadband laser gun sound.

Because the response pattern for this task was inverted and responses were only required on target trials, detection of oddball targets in this task is trivial, leading to overall high performance. Therefore, sustained attention performance in this dataset was quantified using the mean run reaction time (RT) variability which has previously been shown to be robustly related to sustained attention performance in both healthy adults and in populations characterized by sustained attention deficits (Chidharom & Carlisle, 2021; Esterman et al., 2013; Karamacoska et al., 2018; Robertson et al., 1996; Seli et al., 2011; Tamm et al., 2012). Importantly, this measure provides more variability across participants than a measure of sensitivity on a task where performance is at ceiling, as in the current dataset. RT variability is predictive of sustained attention ability such that individuals with more variable pressing show worse performance on sustained attention tasks. Since RT variability has previously been shown to be negatively related to sustained attention performance, we report the inverse of RT variability (mean RT / standard deviation) for ease of comparison with Datasets 1 and 2 in the current study.

### fMRI preprocessing procedure

Functional MRI data for the three datasets underwent the same preprocessing steps in AFNI (Cox, 1996). Preprocessing included the following steps: Removal of leading TRs as previously noted for individual datasets; alignment of functional data to MNI space; regression of covariates of no interest, including a 24-parameter head motion model (6 motion parameters, 6 temporal derivatives, and their squares), mean signal from subject-level white matter and ventricle masks, and mean whole-brain signal; and censoring of volumes for which the derivative of motion parameters exceeded .25 mm or for which more than 10% of the brain were outliers.

### Exclusion criteria

To ensure high-quality data, individual runs were excluded if they did not meet the following criteria regarding head motion inside the scanner and behavioral performance. Runs were excluded if mean framewise head displacement after motion censoring exceeded .15mm, if the maximum head displacement exceeded 4mm, or if greater than 50% of frames were censored during preprocessing. Runs in Datasets 1 and 2 were also excluded if hit rates were more than 2.5 standard deviations below the mean hit rate value. The tasks used in these datasets asked participants to respond to frequent trials (90%), such that good performance would require presses to the vast majority of trials. Therefore, low hit rates for these tasks indicate participant non-compliance. Finally, we excluded runs if behavioral performance, quantified as sensitivity (*d’*) in Datasets 1 and 2 and inverse RT variability in Dataset 3, was greater than 2.5 standard deviations below the mean across all runs within a dataset.

In Dataset 1, two visual runs were removed based on head motion criteria and 6 visual runs were excluded for extremely low hit rates. No auditory runs were removed based on any of the listed criteria. In Dataset 2, 56 participants completed the visual *avCPT* and 55 participants completed the auditory *avCPT*. 9 visual runs and 10 auditory runs were excluded based on head motion criteria. An additional two visual runs and one auditory run were removed due to low hit rates. In the final sample for Dataset 2, 36 participants completed both a visual and an auditory run. For Dataset 3, 47 visual and 44 auditory runs were successfully preprocessed. Preprocessing failed for the remaining 4 visual and 7 auditory runs due to the number of time points censored. No additional runs were removed based on head motion criteria. No runs in any dataset were excluded on the basis of low sensitivity or RT variability measures. The final sample sizes for each dataset and run type were as follows: Dataset 1 included 50 visual and 58 auditory runs, Dataset 2 included 45 visual and 44 auditory runs, and Dataset 3 included 47 visual and 44 auditory runs.

### External validation of sustained attention CPM

Functional MRI data were parcellated into 268 functionally-defined regions of interest (ROIs, Shen et al., 2013). Whole-brain functional connectivity matrices were calculated by correlating the blood oxygen level dependent (BOLD) time courses for a given task run between all pairs of ROIs. Edges in this 268 by 268 matrix provide an index of coactivation similarity between all pairs of regions in the brain for each run.

Our first question of interest was whether a predefined network trained to predict sustained attention performance in a visual task generalized to the present datasets which include both visual and auditory sustained attention tasks. The network tested was defined using connectome-based predictive modeling (CPM; Finn et al., 2015; Rosenberg et al., 2016a; Shen et al., 2017) which identifies a set of edges whose coactivation strength is related to a performance metric across a set of participants. In CPM, the strength of every edge in a functional connectivity matrix is correlated with a behavior of interest, in this case sustained attention performance. The predefined network, referred to in the current manuscript as the *saCPM* (sustained attention CPM) consists of a set of edges whose strength was either positively (757 edges) or negatively (630 edges) correlated with visual *gradCPT* performance across an independent set of participants (*N*=25). Significant edges were defined as those whose network strength was significantly correlated (Pearson’s *r*; p<.01) with visual gradCPT sensitivity (*d’*) across participants. Positively correlated edges are connections whose strength increased with higher sustained attention performance across participants, whereas negatively correlated edges are connections whose strength increased with worse performance. This network is described in previous work by Rosenberg et al., (2016a; 2020) and is shared publicly (https://github.com/monicadrosenberg/Rosenberg_PNAS2020).

Here, we tested whether strength in this predefined network also predicted sustained attention performance in datasets that include novel participants, multiple perceptual modalities, and new behavioral measures of interest. Network strength is defined as the difference between mean connectivity in the high-attention and mean connectivity in the low-attention network for each run in the current datasets. Because strength in the high-attention and low-attention networks will be negatively correlated by nature of how the networks were identified, taking the difference provides a single summary measure which is interpretable. CPM-predicted behavior is a linear transformation of network strength (predicted behavior = *m*\*network strength + *b*, where *m* and *b* are learned during model training). Therefore, for external model validation as we perform in the current set of analyses, the correlation between network strength and observed behavior is mathematically equivalent to correlation between predicted and observed behavioral scores. Network strength values were normalized across participants within dataset for comparison with other analyses. We then tested whether that network strength was related to behavioral performance by calculating the partial Spearman’s *rho* value between network strength and the behavioral measure of interest for visual and auditory runs separately, controlling for mean head motion (mean framewise displacement) in the scanner. Spearman’s rho values were used to mitigate any potential effects of outliers on predictions. However, results are consistent when using Pearson’s correlation.

As a note, we do not apply multiple comparison correction for the present study because all tests of model generalization tested a non-omnibus hypothesis, i.e., that network strength in the trained model will predict sustained attention performance in an independent sample (Garcia-Perez, 2023). Each external validation of model prediction tests a single outcome (significance of correlation between network strength and performance) and therefore multiple comparisons corrections would create unnecessarily large barriers to generalization.

### Modality-specific model construction

Next, we tested whether a network that is trained on fMRI data collected during a sustained attention task performed in a given perceptual modality better predicts performance on a task performed in the same vs. a different modality. To test this, we defined new models on the functional connectivity matrices and behavior in Dataset 1 using a CPM approach (Finn et al., 2015; Rosenberg et al., 2016a; Shen et al., 2017). CPM identifies a set of edges that is correlated, either negatively or positively (Pearson’s *r*, p<.01) with behavioral performance across the training set. For the current analyses, the training set was all visual or auditory runs in Dataset 1. For each edge in a functional connectivity matrix, a Pearson’s correlation is calculated between edge strength and sustained attention performance across the dataset. This is repeated for all edges in the functional connectivity matrix and significant edges are those whose correlation with sustained attention is stronger than a given threshold, in this case, p<.01. Positive network edges are those where connectivity strength is positively related to behavior across an entire training sample, while negative network edges are those whose connectivity is negatively related to behavior across the sample. Significant edges are isolated to represent a network of edges for which edge strength is related to sustained attention in a given dataset. This results in binary edge “masks” consisting of 0s and 1s for both positive and negative networks. Edge masks are used to calculate network strength in independent datasets by calculating the dot product between the binary mask and each individual’s FC matrix and taking the difference between average connectivity strengths in the positive and negative edge networks. Networks were defined on visual and auditory runs of Dataset 1 separately. We then tested the generalizability of these networks by calculating the partial Spearman’s correlation between modality-specific network strength and performance in the left-out datasets 2 and 3 visual and auditory runs, controlling for in-scanner head motion. For these external validation analyses, the correlation between network strength and observed behavioral scores is again equivalent to the correlation between predicted and observed behavioral scores.

To investigate the composition of visual, auditory, and overlapping sustained attention networks, we quantified the relative contribution of canonical brain networks (Finn et al., 2015) to these networks. This functionally-defined canonical network parcellation includes visual networks labeled based on their similarity to resting-state visual networks. There is no comparable auditory network included in this parcellation. However, connections from auditory cortex may be best encompassed by medial frontal and motor networks. We quantified relative contribution to high and low sustained attention networks by calculating the difference between the number of edges identified by high and low networks within and between canonical networks. This relative contribution was normalized by the total number of edges contained in a network. Significance of network contributions was calculated non-parametrically by shuffling edges in the high and low attention networks separately and recalculating the difference in network contribution 1000 times.

We quantified the significance of overlap between our new visual and auditory sustained attention networks using a hypergeometric cumulative distribution function, which calculates the probability of observing the number of overlapping edges given a random sampling with no replacement of two networks of the sizes observed (Rosenberg et al., 2016b). Significance values were calculated in MATLAB as p=1-*hygcdf(x,M,K,N)* where *x* is the number of shared edges between networks of interest, *M* is the total number of functional edges in the matrix (35,778), and *K* and *N* are the number of functional edges the networks of interest.

We tested whether model generalization was biased towards the perceptual modality of training by calculating a measure of modality-specificity for each external validation dataset. Modality-specificity of visual and auditory networks was calculated as the prediction (partial Spearman’s rho) of within-modality generalization (e.g., visual performance predicted by the visual CPM) minus cross-modality generalization (e.g., visual performance predicted by the auditory CPM) for each dataset and modality. We determined significance with a permutation test whereby predicted performance values were shuffled and correlated with observed performance, controlling for head motion. Auditory and visual predicted performance values were shuffled independently and the difference between these partial Spearman’s rho values was calculated. This process was repeated across 5000 iterations to obtain a null distribution of permuted difference scores.

Finally, we tested the contribution of network components to the generalizability of auditory and visual networks. To do so, we calculated whether network strength in reliably predictive edges, i.e., edges that appeared in both visual and auditory predictive networks, was related to sustained attention performance in independent datasets. We hypothesized that these edges would reflect connectivity involved in supramodal sustained attention and therefore would generalize to predict performance in both modalities. We also tested whether edges that appeared only in the visual network or the auditory network would show specificity to their training modality. To do this we calculated network strength in edges that appeared either in the visual network or the auditory network, but not in both. We calculated the modality specificity of visual-only and auditory-only network edges by comparing predictions within and across modality, as described in the previous paragraph.

All preprocessed data and analysis code required to recreate the described analyses are publicly available at https://osf.io/bt2xy/.

## Results

### A predefined visual network generalizes across datasets and modalities

We first tested whether the *saCPM*, a network trained to predict performance on a visual sustained attention task generalized to the current datasets. Sustained attention performance, measured as sensitivity (*d’*) in Dataset 1, ranged from 1.14 to 5.11 in visual runs (M=3.11, SD=.957) and .306 to 3.69 in auditory runs (M=1.51, SD=.679). In Dataset 2, visual *d’* values ranged from 1.08 to 4.40 (M=3.09, SD=.640) and auditory *d’* values ranged from -.112 to 2.22 (M=.968, SD=.599). Inverse RT variability in Dataset 3 ranged from 3.67 to 10.88 (M=7.39, SD=1.76) in visual runs and from 2.76 to 11.47 (M=5.53, SD=1.98) in auditory runs. Mean visual and auditory sustained attention measures were positively related across subjects in all datasets (Spearman’s *rho_1_*=.497, p_1_=9.84*10^-3^; Spearman’s *rho_2_*=.537, p_2_<.001; Spearman’s *rho_3_*=.589, p_3_=.021). However, performance was not perfectly correlated across modalities, such that not all variance in auditory task performance was explained by performance on the visual task. Therefore, successful generalization of the *saCPM* would require that it rely on features which capture the shared, supramodal variance.

For visual task runs, network strength in the *saCPM* was positively related to performance in all datasets (partial *rho_1_*=.230, p_1_=.112; partial *rho_2_*=.317, p_2_=.036; partial *rho_3_*=.343, p_1_=.020) and this relationship was significant in Datasets 2 and 3 (**Figure 1**). While the prediction of visual sustained attention was not significant in Dataset 1, the relationship between network strength and observed performance was in the predicted direction and aligns with predictions in other datasets. As a validation that the *saCPM* captures visual sustained attention performance in Dataset 1, we also tested whether network strength in the saCPM predicted inverse RT variability in this dataset. RT variability was significantly correlated with sustained attention performance in visual runs of Dataset 1 (*r=*.649, p<.001) but may capture more meaningful variance in performance in this dataset. *saCPM* Network strength positively predicted inverse RT variability during the visual task (partial *rho*=.313, p=.028). Therefore, we concluded that this previously-validated network generalizes to predict visual sustained attention performance in Dataset 1. When applied to auditory task runs, *saCPM* network strength significantly predicted auditory sustained attention performance in all three datasets (partial *rho_1_*=.344, p_1_=8.74*10^-3^; partial *rho_2_*=.360, p_2_=.018; partial *rho_3_*=.552, p_1_<.001). Successful generalization of the predefined *saCPM* demonstrates that this network captures features of sustained attention that are general across datasets as well as perceptual modalities.

**Figure 1.**
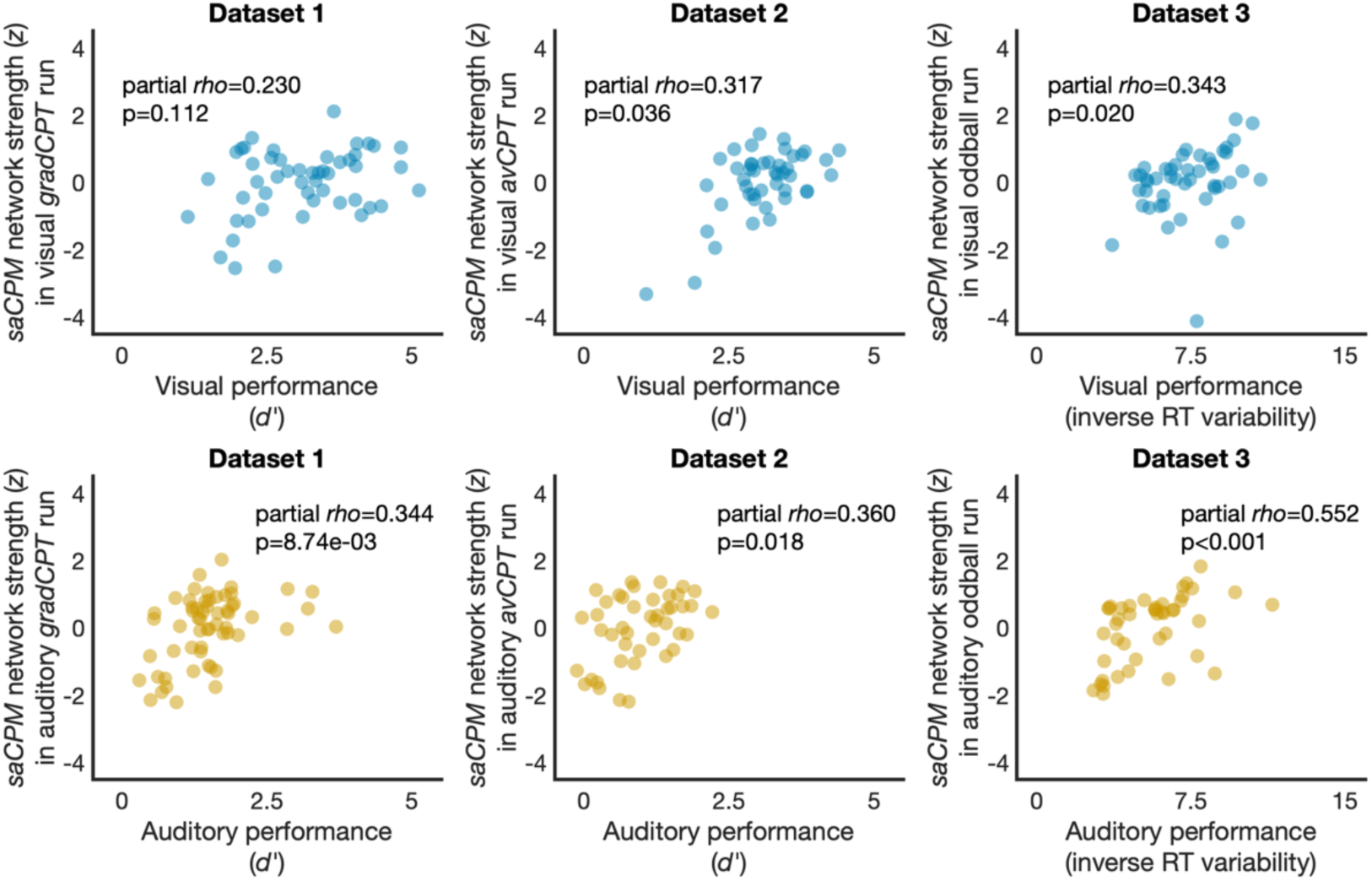
Network strength in the saCPM network significantly predicted visual sustained attention performance in Datasets 2 and 3 and auditory sustained attention performance in all datasets.

### Sustained attention networks are not modality-specific

We next asked whether a model trained on an auditory sustained attention task would generalize to predict performance on other auditory attention tasks better than a model trained on a visual attention task. One option for doing so would be training a new CPM to predict auditory task performance, applying the model to new data, comparing its predictive power to that of the *saCPM*. However, in this scenario any differences in predictive power could be due to differences between training datasets (number of participations, amount and quality of data, etc.) rather than attention modality per se. Thus, to more directly compare the generalizability of auditory and visual attention models, we constructed two new models—an auditory model trained to predict auditory *gradCPT* performance in Dataset 1 and a visual model trained to predict visual *gradCPT* performance in Dataset 1. Dataset 1 was selected as the training dataset because sustained attention performance in this dataset was measured using the *gradCPT*. Thus, networks from this dataset are most comparable to saCPM networks which were trained using the same task. We tested the generalizability of these models within and across perceptual modalities by relating network strength in the visual and auditory networks to visual and auditory sustained attention performance in Datasets 2 and 3, controlling for head motion during task runs. For all results reported below, models were applied to functional connectivity data from an auditory or visual task run and resulting predictions were related to behavioral performance from that same task run.

The visual network generalized to predict visual sustained attention in Dataset 2 (partial *rho*=.329, p=.029) and Dataset 3 (partial *rho*=.305, p=.039; **Figure 2A**). The auditory network similarly generalized to predict auditory sustained attention performance in both datasets (partial *rho*_2_=.376, p_2_=.013; partial *rho_3_*=.537, p_3_<.001). Within-modality generalization confirms that CPM successfully identified networks whose strength predicts out-of-sample sustained attention performance.

**Figure 2.**
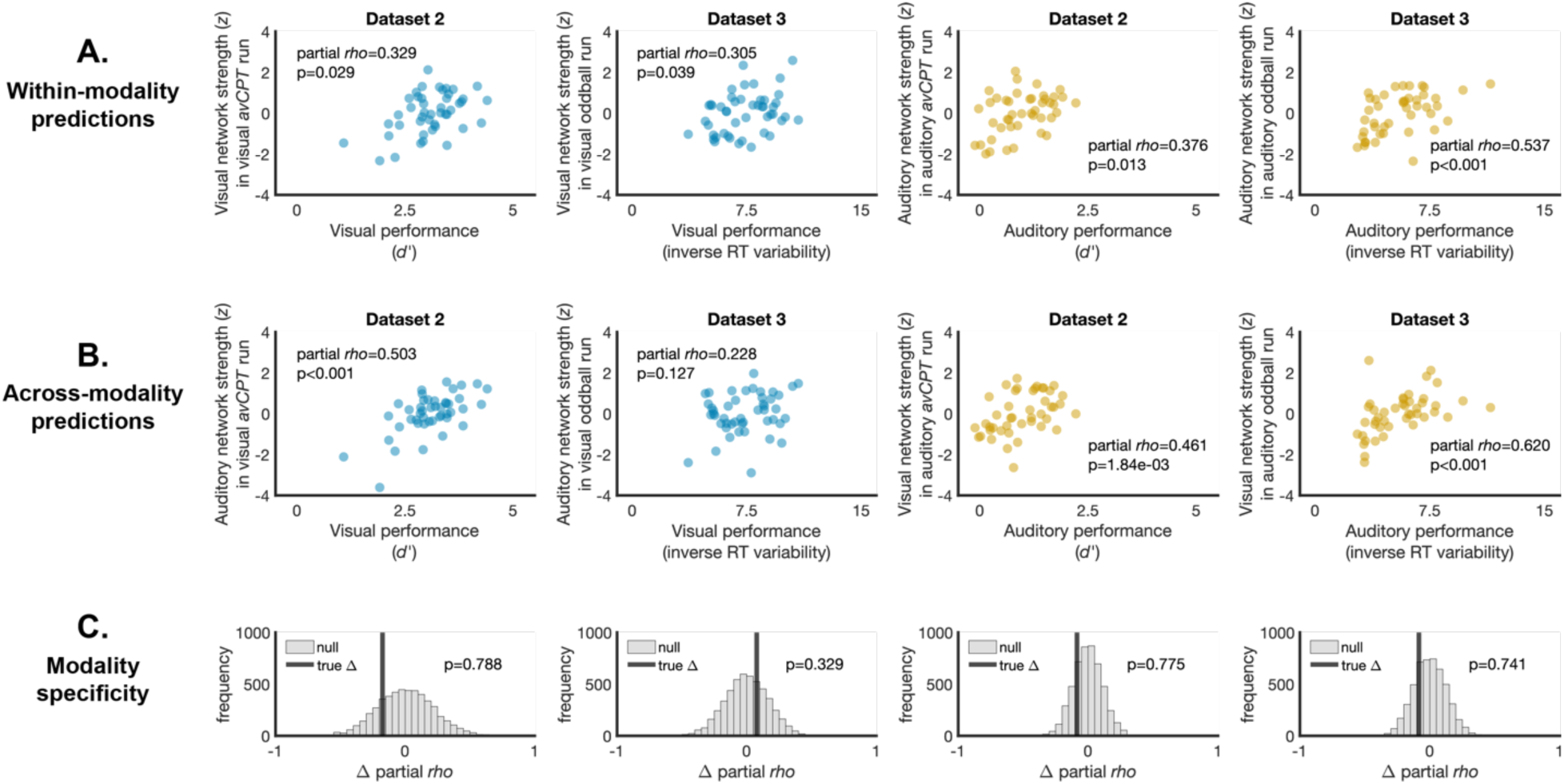
(A) Visual and auditory networks generalized to predict visual and auditory sustained attention performance, respectively, in independent datasets. Network strength is quantified as the difference between the average high and the average low network strength values. (B) The visual network predicted auditory sustained attention in independent datasets and the auditory network predicted visual performance in one dataset. (C) Neither network showed modality-specificity, i.e., generalized better to within-modality prediction than across-modality prediction. The vertical black bar represents the true difference between prediction of task performance from a within-modality model vs. an across-modality model. The gray distribution reflects null differences from predictions of shuffled sustained attention performance. Positive partial *rho* difference values reflect better prediction within vs. across perceptual modality. Negative partial *rho* difference values reflect better performance prediction for a task performed in a different perceptual modality than training.

We next tested whether these networks predicted sustained attention performance when tasks were performed in a different modality. The visual network significantly predicted auditory sustained attention performance in both Dataset 2 (partial *rho*=.461, p=1.84*10^-3^) and Dataset 3 (partial *rho*=.620, p<.001; **Figure 2B**). The auditory network predicted visual sustained attention performance in Dataset 2 (partial *rho*=.503, p<.001) and was positively but not significantly related to visual sustained attention performance in Dataset 3 (partial *rho*=.228, p=.127). Successful generalization across dataset and perceptual modality suggests that sustained attention relies on a modality-general mechanism captured, at least to some extent, by the edges identified by CPM.

To quantify the extent to which networks were modality specific, we calculated the difference between within-modality prediction and across-modality prediction by subtracting the respective partial Spearman’s rho values. We created a null distribution by shuffling model-predicted visual and auditory behavioral performance values independently. The difference between partial Spearman’s rho values was permuted and this was repeated 5000 times. A one-tailed test was used to determine whether the observed difference between within- and between-modality predictions was greater than permuted values. Visual performance was not better predicted by a visual network than an auditory network in Dataset 2 (p=.788) or in Dataset 3 (p=.329; **Figure 2C**). Similarly, auditory performance predictions from the auditory network were not higher than predictions from the visual network in either Dataset 2 (p=.775) or Dataset 3 (p=.741). Across both models, we found no modality-specificity such that the networks identified to predict visual or auditory sustained attention performance did not better predict task performance in the same modality.

We note that the number of runs available in Dataset 1 to train these models differs between auditory (N=58) and visual (N=50) runs. To ensure that training the auditory model on a larger number of runs did not bias the model’s generalizability, we subsampled the number of runs used to train the auditory network to be equal to the number of runs used to train the visual network, i.e., 50 runs. We refit 1000 auditory models using a random subsampling of 50 auditory runs from Dataset 1 and tested how well these models generalized across datasets and modalities. In all cases, prediction from the model trained on the full N=58 sample fell within one standard deviation of the mean prediction from models trained on a smaller sample (Prediction of visual performance: mean partial *rho*_2_=.478, SD_2_=.054; mean partial *rho*_3_=.220, SD_3_=.053; Prediction of auditory performance: mean partial *rho*_2_=.365, SD_2_=.026; mean partial *rho*_3_=.534, SD_3_=.040). Therefore, it is not the case that prediction from the auditory model in the current analyses is biased due to a larger amount of training data.

### Cross-modality generalization is not explained by within-modality performance

Sustained attention performance is reliable across modalities, such that individuals with high visual sustained attention performance tend to have high auditory sustained attention performance. Therefore, cross-modality generalization of network predictions could result simply because cross-modality performance is related to within-modality performance. Another alternative, however, is that network models capture variance above and beyond sustained attention performance consistency. To test this, we included within-modality sustained attention performance as an additional variable in the partial correlation between cross-modality performance and network strength. If models fail to generalize when supramodal sustained attention performance is captured by the additional variable of within-modality performance, this suggests that the generalizability of these models across modalities relies heavily on features related to this shared supramodal ability. If models still generalize after controlling for supramodal sustained attention performance, this would suggest that networks capture unique variance beyond what can be explained by similarity in sustained attention performance across runs.

The partial correlation between saCPM network strength and auditory sustained attention performance remained significant in both Dataset 1 (partial *rho*=.301, p=.024) and Dataset 3 (partial *rho*=.540, p<.001), even when controlling for participants’ visual sustained attention performance. Prediction in Dataset 2 was positive but not significant after controlling for visual sustained attention performance (partial *rho*=.215, p=.183). Therefore, in two of three datasets, generalization of the saCPM to auditory tasks cannot be explained by a correlation between performance across modalities.

We further tested whether generalization of visual and auditory networks trained on Dataset 1 remained after controlling for within-modality performance. Predictions of auditory sustained attention performance from the visual network remained significant after removing variance explained by visual sustained attention performance in Dataset 2 (partial *rho*=.339, p=.033) and Dataset 3 (partial *rho*=.605, p<.001). When controlling for auditory sustained attention performance, predictions of visual sustained attention from auditory networks were significant in Dataset 2 (partial *rho*=.455, p=2.81*10^-3^) and remained non-significant in Dataset 3 (partial *rho*=.096, p=.532). Therefore, generalization of sustained attention networks across task modalities is not simply due to performance similarity across modalities, but rather networks capture sustained attention ability beyond what can be explained by shared supramodal variance.

As a final test of the extent to which cross-modality generalization relies on shared variance in task performance between modalities, we retrained sustained attention networks to predict auditory and visual performance in Dataset 1, controlling for performance in the other modality. In other words, during the feature selection step of visual sustained attention model training, positive and negative network edges were those that were significantly correlated with visual sustained attention performance across individuals in Dataset 1 partialling out auditory sustained attention performance. Similarly, visual sustained attention performance was included as a partial covariate when identifying auditory sustained attention model features. Therefore, these models should no longer capture variance that can be explained by consistency in performance across individuals. A failure of these models to generalize across datasets and modalities would suggest that previous model generalization relied heavily on the shared variance in sustained attention performance across task modality. However, if these models indeed predict sustained attention performance in a modality different than training, it suggests that model features capture relevant variance beyond what can be explained by consistency in performance across modalities.

Cross-modality predictions of visual sustained attention performance from the retrained auditory network were significant in Dataset 2 (partial *rho*=.546, p<.001) and remained non-significant in Dataset 3 (partial *rho*=.092, p=.544). The retrained visual network significantly predicted auditory sustained attention performance in both Dataset 2 (partial *rho*=.379, p=.012) and Dataset 3 (partial *rho*=.494, p<.001). The strength of prediction, i.e., partial *rho* values, were reduced in three of these cross-modal generalizations, suggesting that shared variance at least partially contributed to the generalizability of sustained attention networks. However, models’ ability to significantly generalize across task modality after controlling for the shared variance in task performance suggests that these models do not rely only on this supramodal variance.

### Unique features underlie auditory and visual networks

Is successful cross-modal prediction a result of shared network edges between visual and auditory networks? If CPM identified a largely overlapping set of edges related to performance on both visual and auditory tasks, it should follow that predictions would not be modality-specific. However, if auditory and visual networks are independent, generalization across modalities might suggest that sustained attention performance can be captured by a diverse set of features.

The visual network consisted of 581 positive (high attention) edges and 659 negative (low attention) edges. In the auditory network, 626 edges were positively related to auditory sustained attention performance and 970 edges were negatively related to auditory sustained attention performance. Edgewise contributions to individual networks are grouped into lobe and canonical network groupings (Finn et al., 2015) and visualized in **Figure 3**.

**Figure 3.**
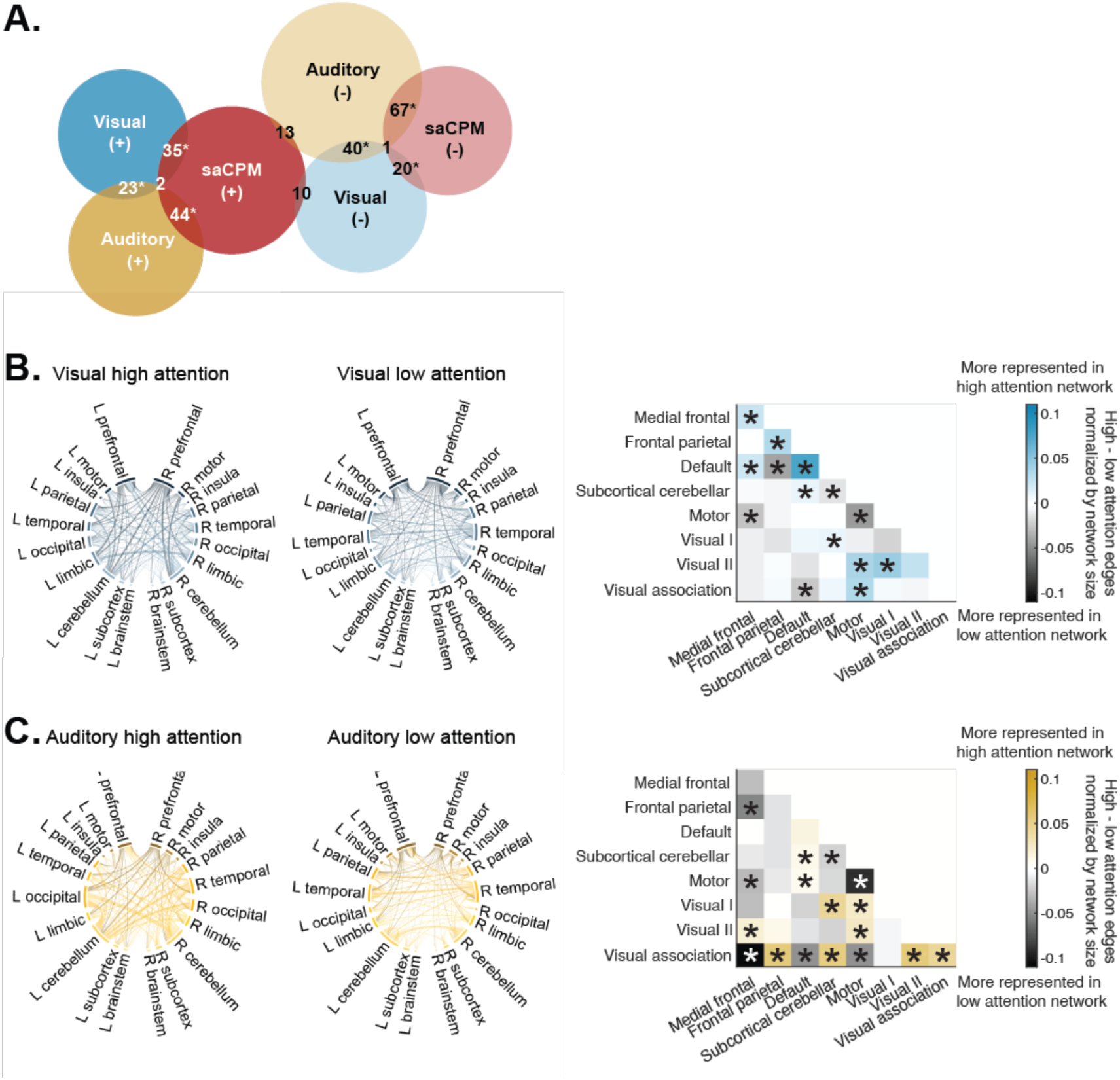
(A) Networks constructed using connectome-based predictive modeling identified shared edges relevant for brain-behavior predictions in both high (+) and low (-) attention networks. Not all overlap between could be visualized in the Venn diagram but is described fully in the text. Stars reflect p<.01. Contributions to network structure grouped by lobe and canonical network differed between (B) visual and (C) auditory networks. Matrices visualize the relative contribution of canonical network edges to high- and low-attention networks. Colors represent the difference between the number of edges in the high and low predictive networks, divided by network size. Stars in the matrix reflect significant contribution to high- or low-attention networks; p<.05, uncorrected. Significance was determined by shuffling high- and low-attention networks and recalculating the contribution of edges to each network 1000 times.

Within-network connections in the medial frontal, frontal parietal, and default mode networks contributed to the visual high-attention network (**Figure 3B**). Connections between the motor network and the visual II, and visual association networks also contributed to the high-attention network, as well as connections between the default mode and medial frontal and subcortical-cerebellar networks. Within-network edges in the motor and subcortical-cerebellar networks contributed to the visual low-attention network. Connections between the default mode and frontal parietal and visual association networks were also stronger in the visual low-attention network, as well as connections between motor and medial frontal networks.

Connections within the visual association network, as well as connections between the visual association and frontal parietal, subcortical-cerebellar, and visual II networks were represented in the auditory high-attention network (**Figure 3C**). Edges shared between motor networks and visual I, visual II, and default mode networks were also represented in the auditory high-attention network. Connections between the subcortical-cerebellar network and visual I and default mode networks contributed significantly to the auditory high-attention network, as well as connections between the medial frontal and visual II networks. Conversely, connections between the visual association network and medial frontal, default mode, and motor networks were strongly represented in the auditory low-attention network. Connections between the medial frontal network and frontal parietal and motor networks were also found more strongly in the auditory low-attention network. Finally, connections within the subcortical-cerebellar networks and motor networks contributed to the auditory low-attention network.

Overlap between visual and auditory networks was significant for both high-attention (25 edges, p<.001) and low-attention networks (41 edges; p<.001; **Figure 3A**). Networks also overlapped with the predefined *saCPM*. Auditory networks significantly overlapped with the *saCPM*, sharing 46 high-attention (p<.001) and 68 low-attention edges (p<.001). The visual network also significantly overlapped with the *saCPM,* sharing 37 high- (p<.001) and 21 low-attention edges (p=3.52*10^-3^). Network overlap in unexpected directions was non-significant in all cases (all ps>.826; **Figure 3A**).

### Predictions from non-overlapping features generalize across modality

Does removing overlapping edges from visual and auditory networks induce modality specificity? It is possible that the generalizability of networks across modality is driven by the subset of shared edges between networks. To ask this, we tested whether edges that were unique to the auditory or visual network—e.g., edges that positively predicted auditory performance but did not predict visual performance—generalized in a modality-specific manner.

Predictions of visual sustained attention performance from visual-unique model edges were significant in Dataset 2 (partial *rho*=.326, p=.031) and positive but non-significant in Dataset 3 (partial *rho*=.277, p=.063). Edges specific to the visual network remained generalizable across perceptual modality such that they predicted auditory sustained attention performance in both Dataset 2 (partial *rho*=.466, p=1.64*10^-3^) and Dataset 3 (partial *rho*=.582, p<.001). We observed no evidence for better prediction for visual sustained attention from a model trained on a visual sustained attention task, even after removing modality-general features (p_2_=.805; p_3_=.365).

Auditory-unique edges significantly predicted auditory sustained attention performance in both Dataset 2 (partial *rho*=.371, p=.014) and Dataset 3 (partial *rho*=.537, p<.001). For visual performance, the auditory-unique network predictions were significant in Dataset 2 (partial *rho*=.511, p<.001) and positive but not significant in Dataset 3 (partial *rho*=.212, p=.158). Again, there was no evidence for modality specificity in predictions from auditory-only network edges (p_2_=.783; p_3_=.606). Therefore, predictions from non-overlapping edges did not result in modality-specific generalization. Instead, even network edges unique to a network trained on one modality captured sustained attention ability in another modality.

### Overlapping features are sufficient for prediction

Within-network edges in the default mode network, as well as edges shared between the default mode and medial frontal networks contributed to the overlapping high-attention network (**Figure 4A**). Additionally, connections shared by the frontal parietal and visual II networks, as well as connections shared between the visual association and subcortical-cerebellar networks were strongly represented in the overlapping high-attention network. This suggests that stronger connections between these networks are associated with higher modality-general sustained attention performance. Conversely, within-network edges in the visual association, subcortical-cerebellar, and motor networks contributed strongly to the overlapping low attention network. These results suggest that strong with-network connectivity in these networks is associated with worse sustained attention.

**Figure 4.**
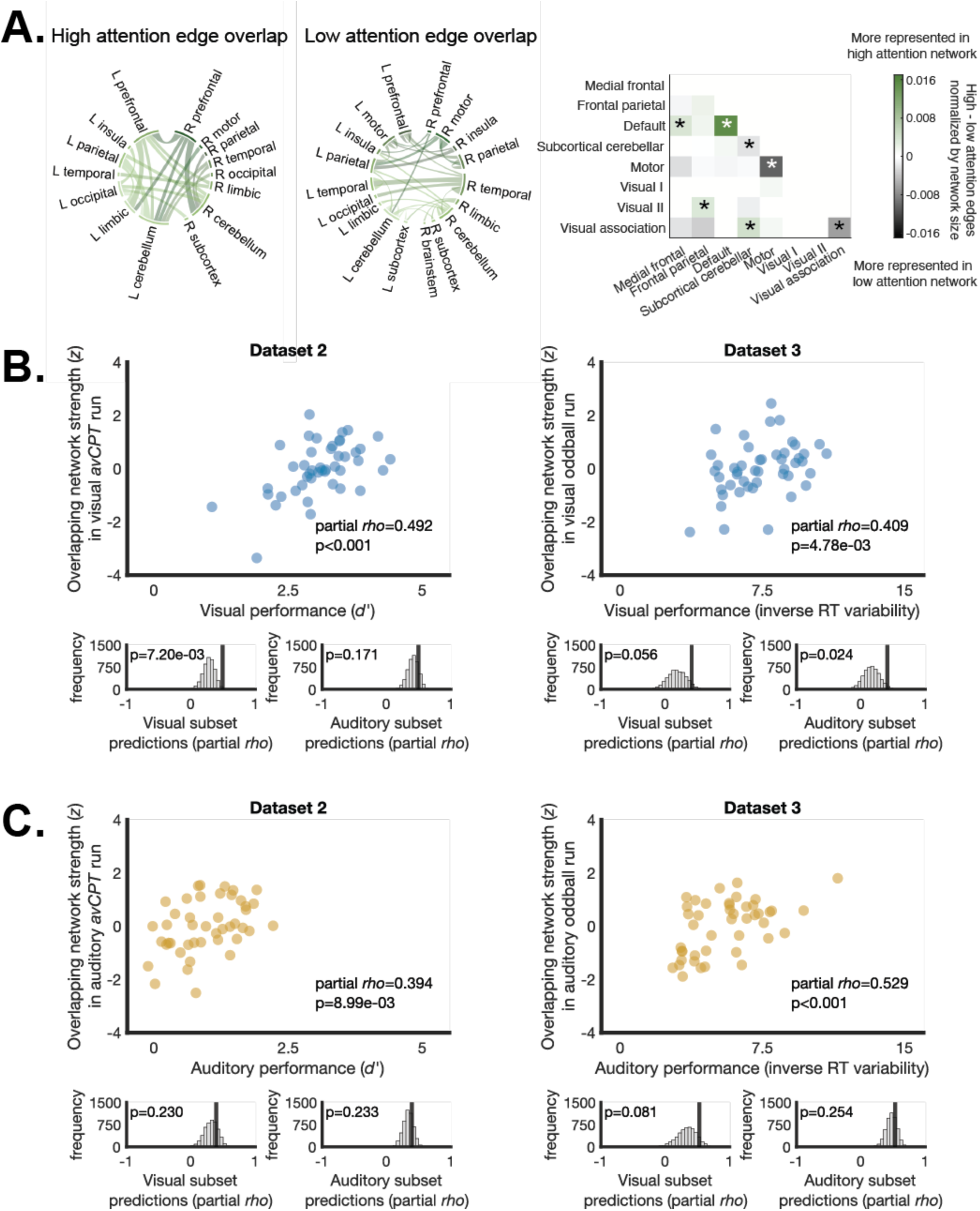
(A) Edges shared by both auditory and visual models trained on Dataset 1 are visualized by lobe. The matrix depicts relative contribution to high and low overlapping predictive networks, grouped into 8 canonical networks. Significance stars on the matrix reflect greater representation of network edges than chance; p<.05, uncorrected. Significance was determined by shuffling network edges 1000 times and recalculating relative contribution to high- and low-attention networks. Network strength in edges shared by the auditory and visual networks predicts (B) visual and (C) auditory sustained attention performance in independent datasets. Histograms beneath each plot depict the extent to which prediction from network strength in the overlapping edges outperforms predictions from equally-sized subsets of edges drawn from either the visual or auditory networks alone (5000 permutations).

We wondered whether the edges shared by both visual and auditory networks defined in Dataset 1 were sufficient to predict visual and auditory sustained attention performance in independent datasets. To ask this, we tested whether strength in the edges shared by high visual and auditory attention networks (25 edges) and low visual and auditory attention networks (41 edges) was related to observed behavioral performance. Results are visualized in **Figure 4B-C**.

We observed robust prediction from these overlapping edges such that network strength in this subset of edges significantly predicted visual performance in Dataset 2 (partial *rho*=.492, p<.001) and Dataset 3 (partial *rho*=.409, p=4.78*10^-3^), as well as auditory sustained attention performance in Dataset 2 (partial *rho*=.394, p=8.99*10^-3^) and Dataset 3 (partial *rho*=.529, p<.001). Therefore, while the number of shared network features was small between visual and auditory networks, these shared features were sufficient for generalizable prediction of sustained attention performance.

Is prediction from a small set of edges specific to the shared edges between visual and auditory networks? To ask this question, we compared the predictive performance from overlapping edges to predictions from equal-sized subsets of visual or auditory network edges. 5000 random subsets were drawn from either the visual or auditory network and network strength from these subsets was related to observed performance. Distributions of partial *rho* values from edge subsets were created for visual and auditory networks separately.

In visual runs, edges that overlapped between the visual and auditory networks outperformed visual sustained attention performance prediction from other edge subsets of the same size drawn from the visual network in Dataset 2 (p=7.20*10^-3^) but did not significantly outperform visual network edge subsets in Dataset 3 (p=.056). Predictions for visual sustained attention performance from overlapping edges did not outperform edge subsets drawn from the auditory network in Dataset 2 (p=.171) but overlap predictions did outperform auditory subset predictions and Dataset 3 (p=.024). This suggests that overlapping visual and auditory predictive edges may carry unique predictive ability related to visual sustained attention performance. When predicting auditory sustained attention performance, predictions from overlapping edges were not better than predictions from edge subsets drawn from either the visual (p_2_=.230; p_3_=.081) or the auditory network (p_2_=.233; p_3_=.254). Therefore, reliable edges provided specific predictive boost in visual runs only.

## Discussion

Prior predictions from connectome-based models of sustained attention may have been more limited than previously thought if they were driven by visual task performance specifically. Here, we tested the extent to which functional networks of sustained attention are modality-specific, with two likely outcomes. First, functional networks or a subset of functional networks may have predicted sustained attention in a modality-specific manner, generalizing better to tasks performed in the same perceptual modality as training. Alternatively, functional networks may not show modality-specificity and predict sustained attention performance for tasks performed in different modalities similarly. Results show evidence for the latter, demonstrating wide-spread cross-modality generalization even when predictive model features are largely unique. This suggests that sustained attention performance can be captured by distributed, supramodal connections in the brain. Further, we demonstrate that both shared and unique edges in visual and auditory networks predict sustained attention performance across modality, showing that both reliable (overlapping) and unreliable (unique) model features can capture relevant brain-behavior relationships.

Work investigating brain-behavior relationships emphasizes that testing model generalizability, and in particular generalizability to external datasets, is the gold standard for the construction of accurate predictive models (Poldrack et al., 2020; Rosenberg & Finn, 2022; Scheinost et al., 2019). Connectome-based predictive models have previously demonstrated robust generalizability to predict relevant cognitive phenotypes across independent samples (Avery et al., 2020; Fountain-Zaragoza et al., 2019; Gao et al., 2020; Kardan et al., 2022; Rosenberg et al., 2016a, 2018, 2020). Therefore, CPM meets this high benchmark for model validity and holds promise for identifying robust and interpretable predictors of cognitive variation.

Here, we test whether CPM-derived functional networks capture variability in sustained attention performance across participants in three independent datasets. All three datasets included fMRI tasks which required sustained attention to stimuli presented either in the visual or auditory domain. However, tasks differed across datasets in several ways, including frequency of responding, selection demands, and inhibitory control. Therefore, successful prediction of performance in these datasets suggests that functional networks successfully capture a signal of sustained attention which is general across all three task contexts rather than a distinct process idiosyncratic to a subset.

We first validate that a network of sustained attention previously defined using a CPM approach, the *saCPM*, generalizes to predict sustained attention performance in these datasets. Previous work has demonstrated that the *saCPM* captures patterns of connectivity related to attention by predicting out-of-sample performance on multiple attention tasks (Fountain-Zaragoza et al., 2019; Kardan et al., 2022; Rosenberg et al., 2018, 2020; Yoo et al., 2022) as well as ADHD symptomatology (Rosenberg et al., 2016a) and variability in narrative engagement within individuals (Song & Rosenberg, 2021). A previous study also found that network strength in the *saCPM* during rest predicted performance on an auditory sustained attention task (Wu et al., 2020). Our results show that the *saCPM* also generalizes to predict sustained attention across perceptual modalities from task connectivity, demonstrating that it captures domain-general signatures of attentional ability. Whereas previous work speculated that selective generalization of the *saCPM* to audiovisual movie engagement, but not audio-only story engagement, was due to modality-specificity of the model (Song & Rosenberg, 2021), our results find no modality bias when predicting individual differences in visual and auditory sustained attention. Rather, the differences in prediction observed in previous work may instead reflect other differences between stimuli, for example in the overall engagement with the narratives.

We further show that models trained on sustained attention tasks performed in separate visual and auditory modalities generalize to predict sustained attention performance both in external datasets and when tasks were performed in a different perceptual modality than in training. This suggests that connectome-based predictive modeling identifies edges that capture variability in sustained attention performance that is not specific to the perceptual modality of the task. These results support previous findings that the ability to sustain attention to visual and auditory information relies to some extent on shared neural mechanisms. We used a CPM approach to identify a subset of edges that significantly predicted both auditory and visual sustained attention performance across individuals in a dataset. This subset of edges predicted both visual and auditory sustained attention performance in independent datasets. Therefore, this overlapping network of edges provides one mechanism that may support a modality-general ability to sustain attention over time.

Importantly, we show that successful generalization across modalities is not simply a mathematical inevitability due to correlations between sustained attention performance across modalities. While performance was reliable across participants regardless of task modality, generalization across modality persisted after controlling for performance in the other task modality during both model training and model testing. Therefore, predictive edges identified by CPM were able to capture relevant variance in sustained attention beyond consistency in performance.

Intriguingly, we observed significant prediction both from overlapping visual and auditory edges as well as modality-specific edges identified using a CPM approach. Therefore, feature reliability, or the identification of the same model features across training sets, was not necessary for successful generalization. These results highlight a distinction between model feature reliability and the ability to predict behavioral phenotypes in an external sample. Previous work has noted this difference, demonstrating that predictive accuracy is not necessarily a result of reliable features (Kragel et al., 2021; Noble et al., 2017; Tian and Zalesky, 2021, although see Chen et al., 2022). Researchers have suggested that a lack of reliability may be a function of the scale at which features are identified, leading to high numbers of model features (Srivastava et al., 2022; Tian & Zalesky, 2021). Here, model features were identified from whole-brain patterns of functional connectivity, consisting of >35,000 pairwise connections between regions. Therefore, it is difficult to determine whether the failure of an edge to be significantly related to performance in both visual and auditory networks is the result of the modality-specificity of the edges or a result of the relatively small scale at which features were identified. As a result, edges identified only in one training set may capture modality-general sustained attention, leading to the significant prediction across modalities observed in the current study.

Individual edge contributions to auditory and visual networks from canonical functional networks varied. We found that connectivity within the default mode network was represented in the auditory high-attention and overlapping high-attention networks but did not significantly contribute to visual predictive networks. Much previous work has related relative increases in default mode network activation with in-the-zone attentional states (Esterman et al., 2013, 2014; Jones et al., 2024; Kucyi et al., 2016, 2017; Fortenbaugh et al., 2018; Song et al., 2022), although changes in activity are not functionally equivalent to changes in connectivity. Past work has shown links between greater within-default mode network connectivity and higher attention (Gordon et al., 2012; Kucyi & Davis, 2014). Further, attention-related disorders are characterized by decreased connectivity within the default mode network (Castellanos et al., 2008; Fair et al., 2010). However, other work has found an inverse or no relationship (Kucyi et al., 2017; Mittner et al. 2014; Esterman et al., 2013), suggesting associations of within-network connectivity of the default mode network with sustained attention are complex. The current findings suggest that stronger within-default mode network connections are associated with higher modality-general sustained attention performance.

We observed a large contribution of within-network edges from the subcortical-cerebellar and motor networks to visual and auditory low-attention networks, as well as the overlapping low-attention network. This is in line with previous work which has implicated greater within-subcortical-cerebellar connectivity in lower sustained attention performance (Fong et al., 2019; Jones et al., 2024; Rosenberg et al., 2016a). Increased within-motor connectivity has similarly been related to poor sustained attention in adolescents, whereas connections between motor and visual regions are increased with better sustained attention (O’Halloran et al., 2018). We observed a similar pattern of results, with connectivity between motor and visual II networks contributing to visual and auditory high-attention networks. We do not see a significant contribution of motor to visual II connectivity to the overlapping high-attention network, suggesting the individual edges may differ between visual and auditory networks. Since the *gradCPT* used to train networks in the current study requires a motor (button press) response, it is possible that connections within and between the motor network are more strongly represented in these networks than would be expected if a different sustained attention task were used for network training. Future work may seek to test the extent to which task demands influence network architecture.

We should also note a few limitations of the current study. First, our analyses utilized a connectome-based predictive modeling approach which sought to identify connections between brain regions whose strength captured variability in modality unique or modality general sustained attention. However, it is likely that functional relationships in the brain, beyond those at the edge level, may differ between task-modality. While outside the scope of the current manuscript, future work may aim to more fully characterize functional differences between task, for example, at the level of graph-theoretic differences between whole-brain connectivity patterns. An additional limitation is the precision of the current predicted sustained attention performance values. Significant correlations between predicted and observed sustained attention performance suggest that our sustained attention networks capture reliable differences in performance and are therefore useful in understanding neural mechanisms involved in sustained attention. However, the current models leave much variance unexplained, which may result from a number of individual, task, and dataset differences. Work aimed at precise predictions of sustained attention performance may choose to include additional variables in predictive models that better-capture this remaining variability.

While the current analyses focused on the generalization of sustained attention networks, a similar question could be asked of predictive networks trained on any cognitive process that can be performed in separate perceptual modalities. For example, it is an open question whether a network trained to predict visual recognition memory across participants would also generalize to predict auditory recognition memory, which is reliably worse (Cohen et al., 2009). Future work testing the validity of brain-based models of cognition should aim to test model generalizability across perceptual modalities to evaluate the extent to which a cognitive process is fully captured by a given model.

Our results demonstrate that functional connectivity-based networks of sustained attention are not specific to the perceptual modality of training, suggesting that these networks capture domain- and modality-general aspects of attention. Both non-overlapping and overlapping, modality-general edges predicted cross-modal sustained attention performance in independent datasets, thereby providing one mechanism by which modality-general sustained attention ability may be supported. These results highlight that the ability to sustain attention to information over time relies on distributed, modality-general connections in the brain and demonstrate the potential for highly-generalizable predictive models constructed from functional connectivity features.

